# Preliminary identification of potential vaccine targets for the COVID-19 coronavirus (SARS-CoV-2) based on SARS-CoV immunological studies

**DOI:** 10.1101/2020.02.03.933226

**Authors:** Syed Faraz Ahmed, Ahmed A. Quadeer, Matthew R. McKay

**Affiliations:** Department of Electronic and Computer Engineering, The Hong Kong University of Science and Technology, Hong Kong, China; Department of Chemical and Biological Engineering, The Hong Kong University of Science and Technology, Hong Kong, China

**Keywords:** Coronavirus, 2019-nCoV, 2019 novel coronavirus, SARS-CoV-2, COVID-19, SARS-CoV, MERS-CoV, T cell epitopes, B cell epitopes, spike protein, nucleocapsid protein, vaccine

## Abstract

The beginning of 2020 has seen the emergence of COVID-19 outbreak caused by a novel coronavirus, Severe Acute Respiratory Syndrome Coronavirus 2 (SARS-CoV-2). There is an imminent need to better understand this new virus and to develop ways to control its spread. In this study, we sought to gain insights for vaccine design against SARS-CoV-2 by considering the high genetic similarity between SARS-CoV-2 and SARS-CoV, which caused the outbreak in 2003, and leveraging existing immunological studies of SARS-CoV. By screening the experimentally-determined SARS-CoV-derived B cell and T cell epitopes in the immunogenic structural proteins of SARS-CoV, we identified a set of B cell and T cell epitopes derived from the spike (S) and nucleocapsid (N) proteins that map identically to SARS-CoV-2 proteins. As no mutation has been observed in these identified epitopes among the available SARS-CoV-2 sequences (as of 9 February 2020), immune targeting of these epitopes may potentially offer protection against this novel virus. For the T cell epitopes, we performed a population coverage analysis of the associated MHC alleles and proposed a set of epitopes that is estimated to provide broad coverage globally, as well as in China. Our findings provide a screened set of epitopes that can help guide experimental efforts towards the development of vaccines against SARS-CoV-2.

## Introduction

The ongoing outbreak of COVID-19 in the Chinese city of Wuhan (Hubei province) (C. Wang, Horby, Hayden, & Gao, 2020) and its alarmingly quick transmission to 25 other countries across the world (Centers-of-Disease-Control-and-Prevention, 2020) resulted in the World Health Organization (WHO) declaring a global health emergency on 30 January 2020 (World-Health-Organization, 2020b). This came just one month after the first reported case on 31 December 2019 (https://www.who.int/emergencies/diseases/novel-coronavirus-2019). WHO, in its first emergency meeting (World-Health-Organization, 2020a), estimated the fatality rate of COVID-19 to be around 4%. Worldwide collaborative efforts from scientists are underway to understand the novel and rapidly spreading virus that causes this disease, SARS-CoV-2 (originally tentatively named 2019-nCoV), and to develop effective interventions for controlling and preventing it (Heymann, 2020; Huang et al., 2020; Xin Liu & Wang, 2020; Zhou et al., 2020).

Coronaviruses are positive-sense single-stranded RNA viruses belonging to the family Coronaviridae. These viruses mostly infect animals, including birds and mammals. In humans, they generally cause mild respiratory infections, such as those observed in the common cold. However, some recent human coronavirus infections have resulted in lethal endemics, which include the SARS (Severe Acute Respiratory Syndrome) and MERS (Middle East Respiratory Syndrome) endemics. Both of these are caused by zoonotic coronaviruses that belong to the genus Betacoronavirus within Coronaviridae. SARS-CoV originated from Southern China and caused an endemic in 2003. A total of 8,098 cases of SARS were reported globally, including 774 associated deaths, and an estimated case-fatality rate of 14-15% (https://www.who.int/csr/sars/archive/2003_05_07a/en/). The first case of MERS occurred in Saudi Arabia in 2012. Since then, a total of 2,494 cases of infection have been reported, including 858 associated deaths, and an estimated high case-fatality rate of 34.4% (https://www.who.int/emergencies/mers-cov/en/). While no case of SARS-CoV infection has been reported since 2004, MERS-CoV has been around since 2012 and has caused multiple sporadic outbreaks in different countries.

Like SARS-CoV and MERS-CoV, the recent SARS-CoV-2 belongs to the Betacoronavirus genus (Lu et al., 2020). It has a genome size of ~30 kilobases which, like other coronaviruses, encodes for multiple structural and non-structural proteins. The structural proteins include the spike (S) protein, the envelope (E) protein, the membrane (M) protein, and the nucleocapsid (N) protein. With SARS-CoV-2 being discovered very recently, there is currently a lack of immunological information available about the virus (e.g., information about immunogenic epitopes eliciting antibody or T cell responses). Preliminary studies suggest that SARS-CoV-2 is quite similar to SARS-CoV based on the full-length genome phylogenetic analysis (Lu et al., 2020; Zhou et al., 2020), and the putatively similar cell entry mechanism and human cell receptor usage (Hoffmann et al., 2020; Letko & Munster, 2020; Zhou et al., 2020). Due to this apparent similarity between the two viruses, previous research that has provided an understanding of protective immune responses against SARS-CoV may potentially be leveraged to aid vaccine development for SARS-CoV-2.

Various reports related to SARS-CoV suggest a protective role of both humoral and cell-mediated immune responses. For the former case, antibody responses generated against the S protein, the most exposed protein of SARS-CoV, have been shown to protect from infection in mouse models (Deming et al., 2006; Graham et al., 2012; Yang et al., 2004). In addition, multiple studies have shown that antibodies generated against the N protein of SARS-CoV, a highly immunogenic and abundantly expressed protein during infection (Lin et al., 2003), were particularly prevalent in SARS-CoV-infected patients (X Liu et al., 2004; J. Wang et al., 2003). While being effective, the antibody response was found to be short-lived in convalescent SARS-CoV patients (Tang et al., 2011). In contrast, T cell responses have been shown to provide long-term protection (Fan et al., 2009; Peng et al., 2006; Tang et al., 2011), even up to 11 years post-infection (Ng et al., 2016), and thus has attracted more interest for a prospective vaccine against SARS-CoV [reviewed in (W. J. Liu et al., 2017)]. Among all SARS-CoV proteins, T cell responses against the structural proteins have been found to be the most immunogenic in peripheral blood mononuclear cells of convalescent SARS-CoV patients as compared to the non-structural proteins (C. K.-F. Li et al., 2008). Further, of the structural proteins, T cell responses against the S and N proteins have been reported to be the most dominant and long-lasting (Channappanavar, Fett, Zhao, Meyerholz, & Perlman, 2014).

Here, by analysing available experimentally-determined SARS-CoV-derived B cell epitopes (both linear and discontinuous) and T cell epitopes, we identify and report those that are completely identical and comprise no mutation in the available SARS-CoV-2 sequences (as of 9 February 2020). These epitopes have the potential, therefore, to elicit a cross-reactive/effective response against SARS-CoV-2. We focused particularly on the epitopes in the S and N structural proteins due to their dominant and long-lasting immune response previously reported against SARS-CoV. For the identified T cell epitopes, we additionally incorporated the information about the associated MHC alleles to provide a list of epitopes that seek to maximize population coverage globally, as well as in China. Our presented results can potentially narrow down the search for potent targets for an effective vaccine against SARS-CoV-2, and help guide experimental studies focused on vaccine development.

## Materials and Methods

### Acquisition and processing of sequence data

A total of 80 whole genome sequences of SARS-CoV-2 were downloaded on 9 February 2020 from the GISAID database (https://www.gisaid.org/CoV2020/) (Table S1). We excluded five sequences (EPI_ISL_ 402120, EPI_ISL_402121, EPI_ISL_402126, EPI_ISL_403928 and EPI_ISL_403931) that likely have spurious mutations resulting from sequencing errors (http://virological.org/t/novel-2019-coronavirus-genome/319/18, http://virological.org/t/clock-and-tmrca-based-on-27-genomes/347, and http://virological.org/t/novel-2019-coronavirus-genome/319/11) and two sequences (EPI_ISL_406959 and EPI_ISL_406960) that were partial genomes. These nucleotide sequences were aligned to the GenBank reference sequence (accession ID: NC_045512.2) and then translated into amino acid residues according to the coding sequence positions provided along the reference sequence for SARS-CoV-2 proteins (orf1a, orf1b, S, ORF3a, E, M, ORF6, ORF7a, ORF7b, ORF8, N, and ORF10). These sequences were aligned separately for each protein using the MAFFT multiple sequence alignment program (Katoh & Standley, 2013). Reference protein sequences for SARS-CoV and MERS-CoV were obtained following the same procedure from GenBank using the accession IDs NC_004718.3 and NC_019843.3, respectively.

### Acquisition and filtering of epitope data

SARS-CoV-derived B cell and T cell epitopes were searched on the NIAID Virus Pathogen Database and Analysis Resource (ViPR)(https://www.viprbrc.org/; accessed 9 February 2020) (Pickett et al., 2012) by querying for the virus species name: “Severe acute respiratory syndrome-related coronavirus” from human hosts. We limited our search to include only the experimentally-determined epitopes that were associated with at least one positive assay: (i) positive B cell assays (e.g., enzyme-linked immunosorbent assay (ELISA)-based qualitative binding) for B cell epitopes; and (ii) either positive T cell assays (such as enzyme-linked immune absorbent spot (ELISPOT) or intracellular cytokine staining (ICS) IFN-γ release), or positive major histocompatibility complex (MHC) binding assays for T cell epitopes. Strictly speaking, the latter set of epitopes, determined using positive MHC binding assays, are candidate epitopes, since a T cell response has not been confirmed experimentally. However, for brevity and to be consistent with the terminology used in the ViPR database, we will not make this qualification, and will simply refer to them as epitopes in this study. The number of B cell and T cell epitopes obtained from the database following the above procedure is listed in Table 1.

**Table 1.** Filtering criteria and corresponding number of SARS-CoV-derived epitopes obtained from the ViPR database.

| Filtering criteria |  | Number of epitopes |
| --- | --- | --- |
| Positive T cell assays | T cell epitopes | 115 |
| Positive MHC binding assays | T cell epitopes | 959 |
| Positive B cell assays | Linear B cell epitopes | 298 |
|  | Discontinuous B cell epitopes | 6 |

### Population-coverage-based T cell epitope selection

Population coverages for sets of T cell epitopes were computed using the tool provided by the Immune Epitope Database (IEDB) (http://tools.iedb.org/population/; accessed 9 February 2020) (Vita et al., 2019). This tool uses the distribution of MHC alleles (with at least 4-digit resolution, e.g., A*02:01) within a defined population (obtained from http://www.allelefrequencies.net/) to estimate the population coverage for a set of T cell epitopes. The estimated population coverage represents the percentage of individuals within the population that are likely to elicit an immune response to at least one T cell epitope from the set. To identify the set of epitopes associated with MHC alleles that would maximize the population coverage, we adopted a greedy approach: (i) we first identified the MHC allele with the highest individual population coverage and initialized the set with their associated epitopes, then (ii) we progressively added epitopes associated with other MHC alleles that resulted in the largest increase of the accumulated population coverage. We stopped when no increase in the accumulated population coverage was observed by adding epitopes associated with any of the remaining MHC alleles.

### Constructing the phylogenetic tree

We used the publicly available software PASTA v1.6.4 [28] to construct a maximum-likelihood phylogenetic tree of each structural protein using the unique set of sequences in the available data of SARS-CoV, MERS-CoV, and SARS-CoV-2. We additionally included the Zaria Bat coronavirus strain (accession ID: HQ166910.1) to serve as an outgroup. The appropriate parameters for tree estimation are automatically selected in the software based on the provided sequence data. For visualizing the constructed phylogenetic trees, we used the publicly available software Dendroscope v3.6.3 [29]. Each constructed tree was rooted with the outgroup Zaria Bat coronavirus strain, and circular phylogram layout was used.

## Results

### Structural proteins of SARS-CoV-2 are genetically similar to SARS-CoV, but not to MERS-CoV

SARS-CoV-2 has been observed to be close to SARS-CoV—much more so than MERS-CoV—based on full-length genome phylogenetic analysis (Lu et al., 2020; Zhou et al., 2020). We checked whether this is also true at the level of the individual structural proteins (S, E, M, and N). A straightforward reference-sequence-based comparison indeed confirmed this, showing that the M, N, and E proteins of SARS-CoV-2 and SARS-CoV have over 90% genetic similarity, while that of the S protein was notably reduced (but still high) (Figure 1a). The similarity between SARS-CoV-2 and MERS-CoV, on the other hand, was substantially lower for all proteins (Figure 1a); a feature that was also evident from the corresponding phylogenetic trees (Figure 1b). We note that while the former analysis (Figure 1a) was based on the reference sequence of each coronavirus, it is indeed a good representative of the virus population, since few amino acid mutations have been observed in the corresponding sequence data (Figure S1). It is also noteworthy that while MERS-CoV is the more recent coronavirus to have infected humans, and is comparatively more recurrent (causing outbreaks in 2012, 2015, and 2018) (https://www.who.int/emergencies/mers-cov/en/), SARS-CoV-2 is closer to SARS-CoV, which has not been observed since 2004.

**Figure 1.**
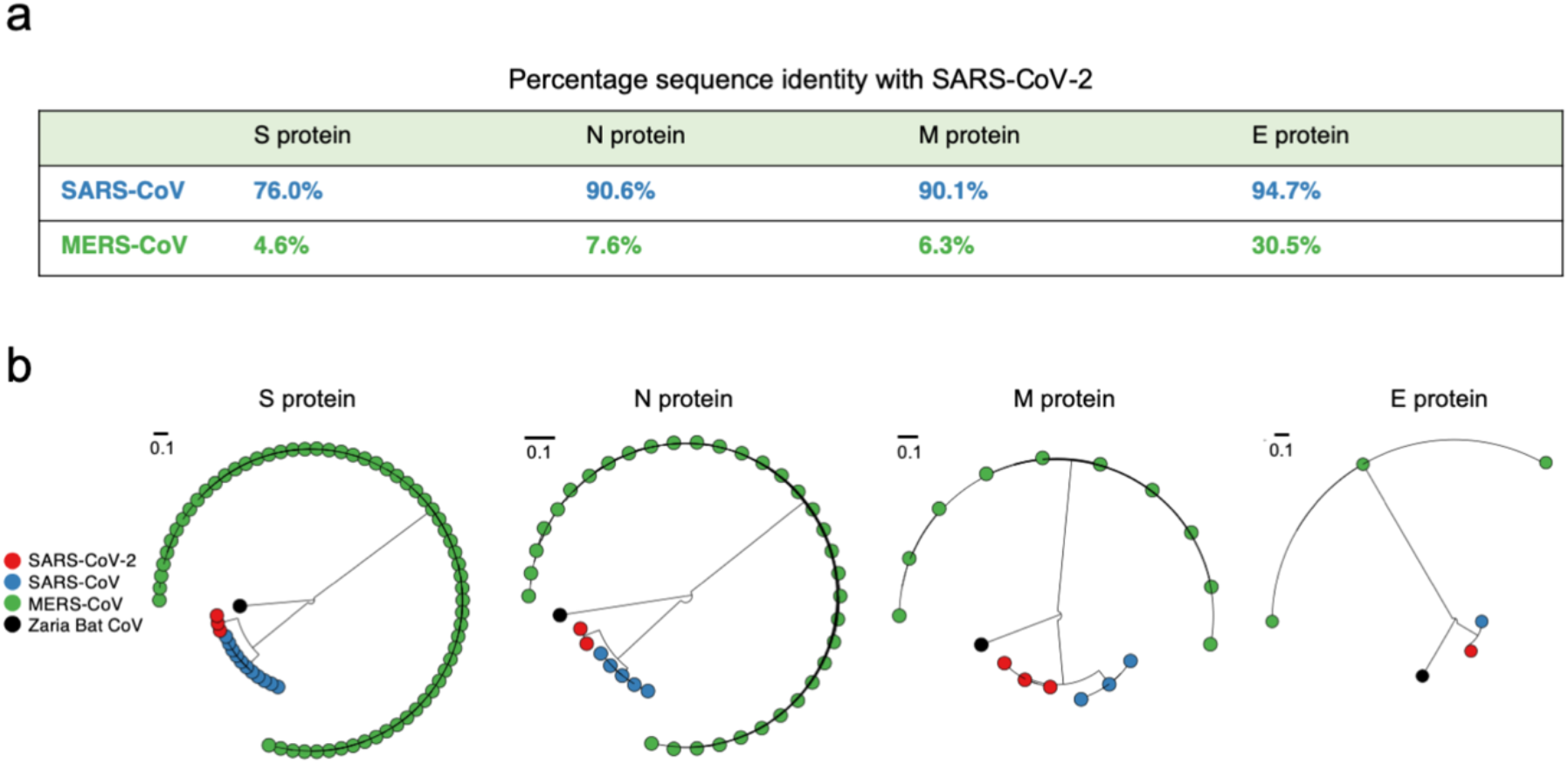
Comparison of the similarity of structural proteins of SARS-CoV-2 with the corresponding proteins of SARS-CoV and MERS-CoV. (a) Percentage genetic similarity of the individual structural proteins of SARS-CoV-2 with those of SARS-CoV and MERS-CoV. The reference sequence of each coronavirus (Materials and Methods) was used to calculate the percentage genetic similarity. (b) Circular phylogram of the phylogenetic trees of the four structural proteins. All trees were constructed based on the available unique sequences using PASTA (Mirarab et al., 2015) and rooted with the outgroup Zaria Bat CoV strain (accession ID: HQ166910.1).

Given the close genetic similarity between the structural proteins of SARS-CoV and SARS-CoV-2, we attempted to leverage immunological studies of the structural proteins of SARS-CoV to potentially aid vaccine development for SARS-CoV-2. We focused specifically on the S and N proteins as these are known to induce potent and long-lived immune responses in SARS-CoV (Channappanavar et al., 2014; Deming et al., 2006; Graham et al., 2012; W. J. Liu et al., 2017; X Liu et al., 2004; J. Wang et al., 2003; Yang et al., 2004). We used the available SARS-CoV-derived experimentally-determined epitope data (see Materials and Methods) and searched to identify T cell and B cell epitopes that were identical— and hence potentially cross-reactive—across SARS-CoV and SARS-CoV-2. We first report the analysis for T cell epitopes, which have been shown to provide a long-lasting immune response against SARS-CoV (Channappanavar et al., 2014), followed by a discussion of B cell epitopes.

### Mapping the SARS-CoV-derived T cell epitopes that are identical in SARS-CoV-2, and determining those with greatest estimated population coverage

The SARS-CoV-derived T cell epitopes used in this study were experimentally-determined from two different types of assays (Pickett et al., 2012): (i) positive T cell assays, which tested for a T cell response against epitopes, and (ii) positive MHC binding assays, which tested for epitope-MHC binding. We aligned these T cell epitopes across the SARS-CoV-2 protein sequences. Among the 115 T cell epitopes that were determined by positive T cell assays (Table 1), we found that 27 epitope-sequences were identical within SARS-CoV-2 proteins and comprised no mutation in the available SARS-CoV-2 sequences (as of 9 February 2020) (Table 2). Interestingly, all of these were present in either the N (16) or S (11) protein. MHC binding assays were performed for 19 of these 27 epitopes, and these were reported to be associated with only five distinct MHC alleles (at 4-digit resolution): HLA-A*02:01, HLA-B*40:01, HLA-DRA*01:01, HLA-DRB1*07:01, and HLA-DRB1*04:01. Consequently, the accumulated population coverage of these epitopes (see Materials and Methods for details) is estimated to not be high for the global population (59.76%), and was quite low for China (32.36%). For the remaining 8 epitopes, since the associated MHC alleles are unknown, they could not be used in the population coverage computation. Additional MHC binding tests to identify the MHC alleles that bind to these 8 epitopes may reveal additional distinct alleles, beyond the five determined so far, that may help to improve population coverage.

**Table 2.**
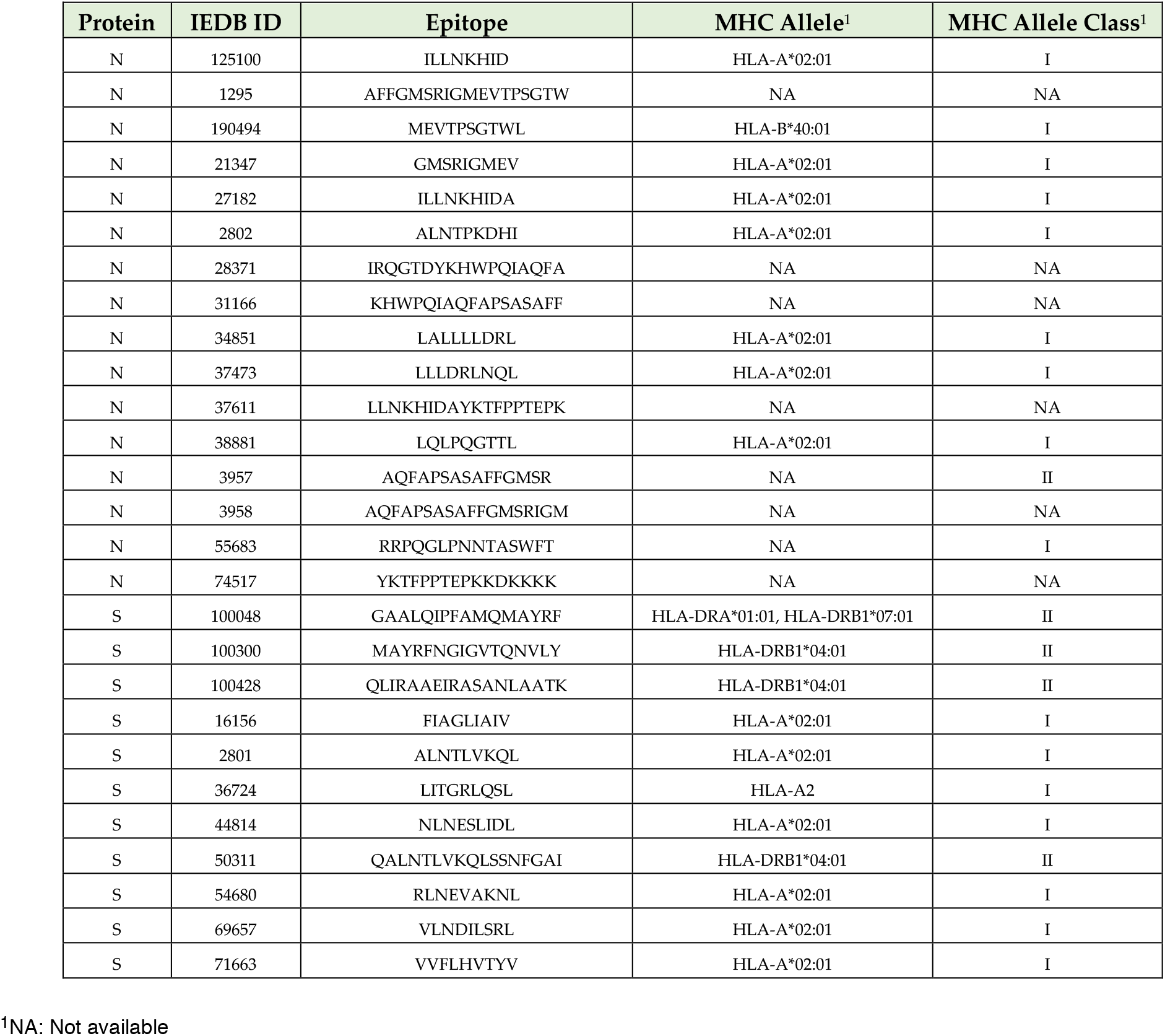
SARS-CoV-derived T cell epitopes obtained using positive T cell assays that are identical in SARS-CoV-2 (27 epitopes in total).

To further expand the search and identify potentially effective T cell targets covering a higher percentage of the population, we next additionally considered the set of T cell epitopes that have been experimentally-determined from positive MHC binding assays (Table 1), but, unlike the previous epitope set, their ability to induce a T cell response against SARS-CoV was not experimentally determined. Nonetheless, they also present promising candidates for inducing a response against SARS-CoV-2. For the expanded set of epitopes, all of which have at least one positive MHC binding assay, we found that 264 epitope-sequences have an identical match in SARS-CoV-2 proteins and have associated MHC allele information available (listed in Table S2). Of these 264 epitopes, ~80% were MHC Class I restricted epitopes (Table S3). Importantly, 102 of the 264 epitopes were derived from either the S (66) or N (36) protein. Mapping all 66 S-derived epitopes onto the resolved crystal structure of the SARS-CoV S protein (Figure 2) revealed that 3 of these (GYQPYRVVVL, QPYRVVVLSF, and PYRVVVLSF) were located entirely in the SARS-CoV receptor-binding motif, known to be important for virus cell entry (F. Li, 2005).

**Figure 2.**
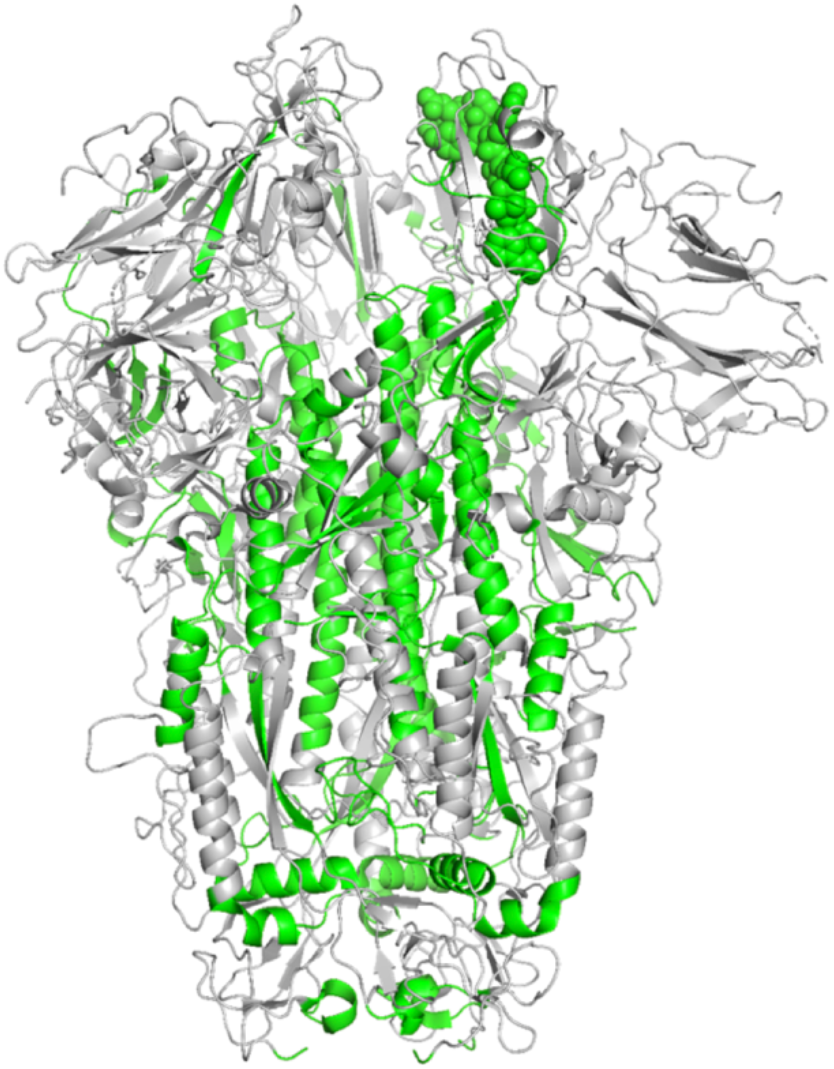
Location of identified T cell epitopes on the SARS-CoV S protein structure (PDB ID: 5XLR). Residues of the SARS-CoV-derived T cell epitopes (determined using positive MHC binding assays and that were identical in SARS-CoV-2) are shown in green color. The 3 overlapping epitopes (GYQPYRVVVL, QPYRVVVLSF, PYRVVVLSF) that lie within the SARS-CoV receptor-binding motif are shown as spheres.

Similar to previous studies on HIV and HCV (Ahmed, Quadeer, Morales-Jimenez, & McKay, 2019; Dahirel et al., 2011; Quadeer et al., 2014), we estimated population coverages for various combinations of MHC alleles associated with these 102 epitopes. Our aim was to determine sets of epitopes associated with MHC alleles with maximum population coverage, potentially aiding the development of subunit vaccines against SARS-CoV-2. For selection, we adopted a greedy computational approach (see Materials and Methods), which identified a set of T cell epitopes estimated to maximize global population coverage. This set comprised of multiple T cell epitopes associated with 20 distinct MHC alleles and was estimated to provide an accumulated population coverage of 96.29% (Table 3). We also computed the population coverage of this specific set of epitopes in China, the country most affected by the COVID-19 outbreak, which was estimated to be slightly lower (88.11%) as certain MHC alleles (e.g., HLA-A*02:01) associated with some of these epitopes are less frequent in the Chinese population (Table 3). Repeating the same greedy approach but focusing on the Chinese population, instead of a global population, the maximum population coverage was estimated to be 92.76% (Table S4).

**Table 3.**
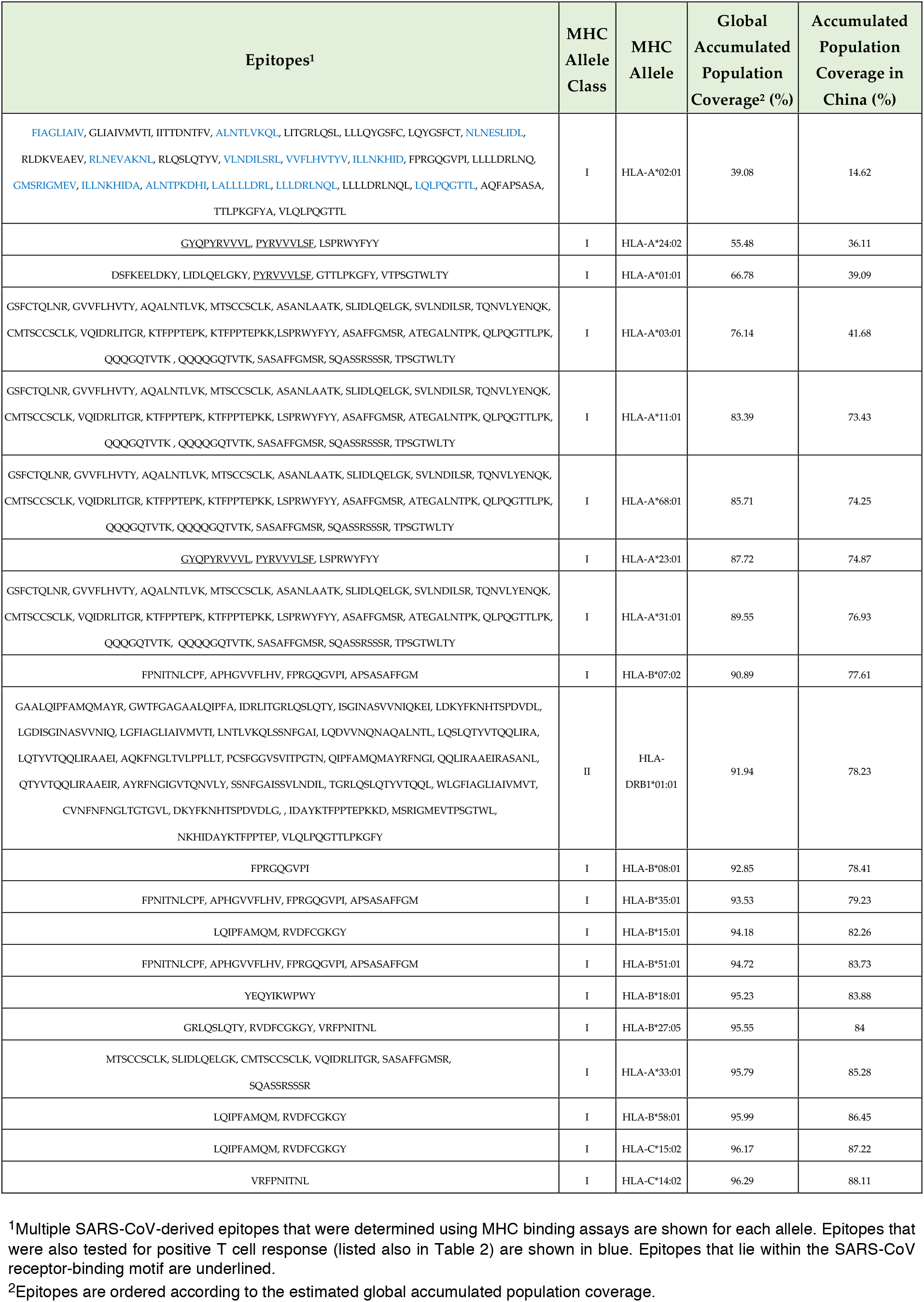
Set of the SARS-CoV-derived S and N protein T cell epitopes (obtained from positive MHC binding assays) that are identical in SARS-CoV-2 and that maximize estimated population coverage globally (87 distinct epitopes).

Due to the promiscuous nature of binding between peptides and MHC alleles, multiple S and N peptides were reported to bind to individual MHC alleles. Thus, while we list all the S and N epitopes that bind to each MHC allele (Table 3), the estimated maximum population coverage may be achieved by selecting at least one epitope for each listed MHC allele. Likewise, many individual S and N epitopes were found to be presented by multiple alleles and thereby estimated to have varying global population coverage (Table S5).

### Mapping the SARS-CoV-derived B cell epitopes that are identical in SARS-CoV-2

Similar to T cell epitopes, we used in our study the SARS-CoV-derived B cell epitopes that have been experimentally-determined from positive B cell assays (Pickett et al., 2012). These epitopes were classified as: (i) linear B cell epitopes (antigenic peptides), and (ii) discontinuous B cell epitopes (conformational epitopes with resolved structural determinants).

We aligned the 298 linear B cell epitopes (Table 1) across the SARS-CoV-2 proteins and found that 50 epitope-sequences, all derived from structural proteins, have an identical match and comprised no mutation in the available SARS-CoV-2 protein sequences (as of 9 February 2020). Interestingly, a large number (45) of these were derived from either the S (23) or N (22) protein (Table 4), while the remaining (5) were from the M and E proteins (Table S6).

**Table 4.**
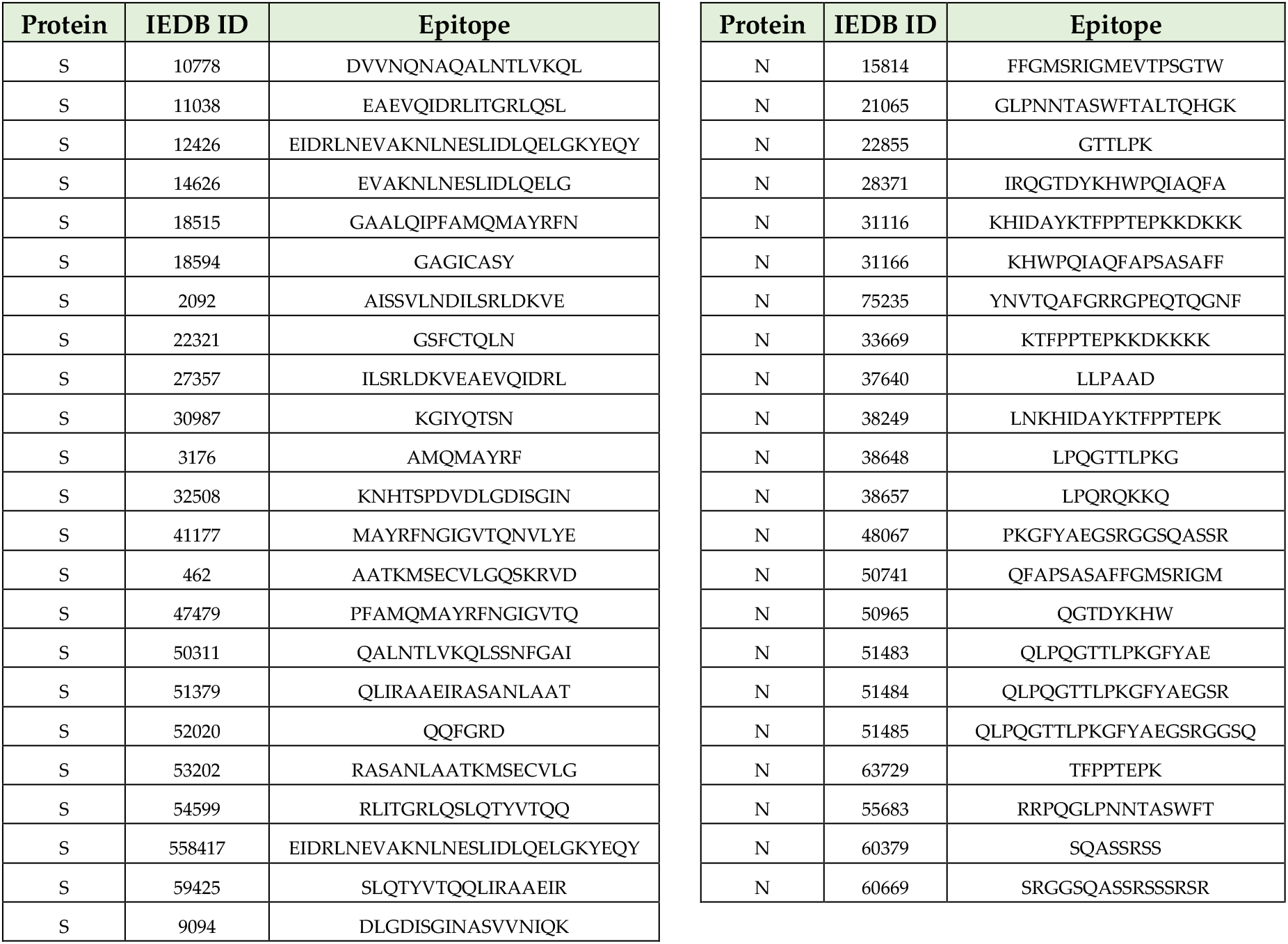
SARS-CoV-derived linear B cell epitopes from S (23) and N (22) proteins that are identical in SARS-CoV-2 (45 epitopes in total).

On the other hand, all 6 SARS-CoV-derived discontinuous B cell epitopes obtained from the ViPR database (Table 5) were derived from the S protein. Based on the pairwise alignment between the SARS-CoV and SARS-CoV-2 reference sequences (Figure S2), we found that none of these mapped identically to the SARS-CoV-2 S protein, in contrast to the linear epitopes. For 3 of these discontinuous B cell epitopes there was a partial mapping, with at least one site having an identical residue at the corresponding site in the SARS-CoV-2 S protein (Table 5).

**Table 5.**
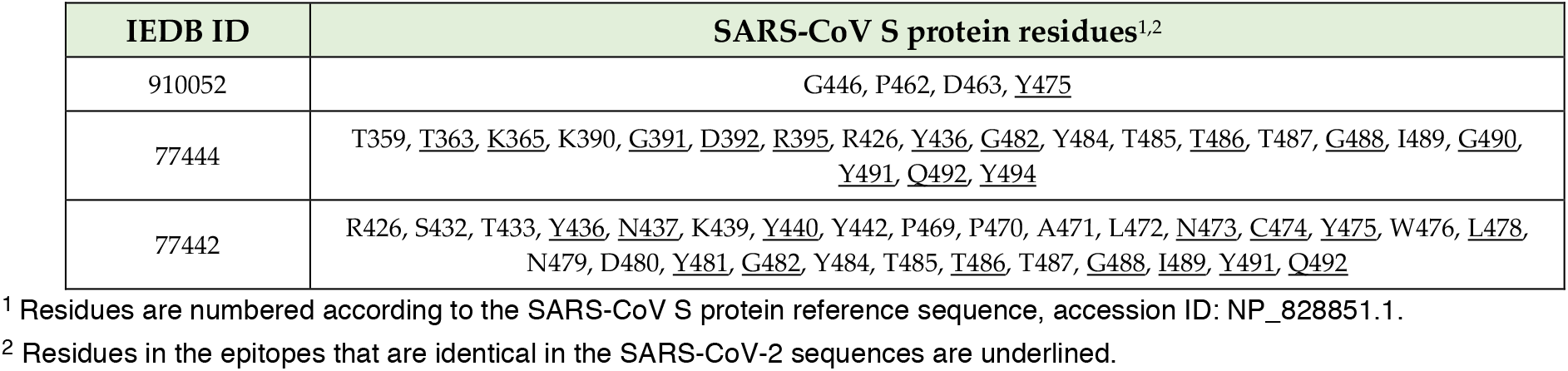
SARS-CoV-derived discontinuous B cell epitopes that have at least one site with an identical amino acid to the corresponding site in SARS-CoV-2.

Mapping the residues of the B cell epitopes onto the available structure of the SARS-CoV S protein revealed that the linear epitopes (Table 4) map to seemingly less-exposed regions, away from the most exposed region, the “spike head” (Figure 3a). In contrast, all discontinuous B cell epitopes (Table 5) map onto the spike head region (Figure 3b, *top panel*), which contains the receptor-binding motif of the SARS-CoV S protein (F. Li, 2005). We observe that few residues of the discontinuous epitopes that lie within the receptor-binding motif are identical within SARS-CoV and SARS-CoV-2 (Figure 3b). It has been reported recently that SARS-CoV-2 is able to bind to the same receptor as SARS-CoV (ACE2) for cell entry, despite having multiple amino acid differences with SARS-CoV’s receptor-binding motif (Hoffmann et al., 2020; Letko & Munster, 2020; Lu et al., 2020; Zhou et al., 2020). Whether the antibodies specific to this motif maintain their binding and elicit an immune response against SARS-CoV-2 warrants further experimental investigation.

**Figure 3.**
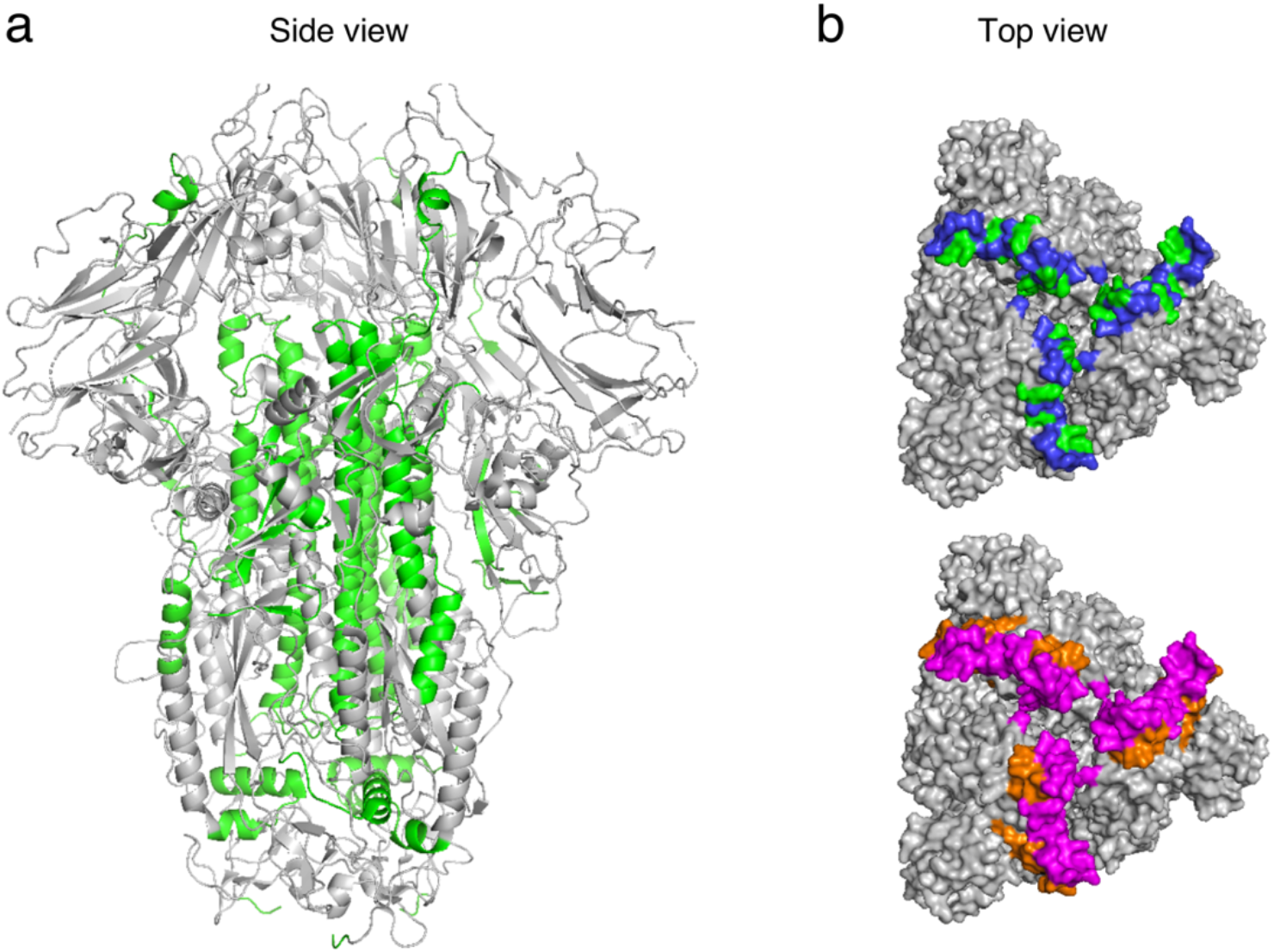
Location of the SARS-CoV-derived B cell epitopes on the SARS-CoV S protein structure (PDB ID: 5XLR). (a) Residues of the linear B cell epitopes, that were identical in SARS-CoV-2 (Table 4), are shown in green color. (b) (*Top panel*) Location of discontinuous B cell epitopes that share at least one identical residue with corresponding SARS-CoV-2 sites (Table 5). Identical epitope residues are shown in green color, while the remaining residues are shown in blue color. (*Bottom panel*) Location of the receptor-binding motif of SARS-CoV. Residues in discontinuous B cell epitopes within the motif are indicated in magenta color, while the remaining motif residues are shown in orange color.

## Discussion

The quest for a vaccine against the novel SARS-CoV-2 is recognized as an urgent problem. Effective vaccination could indeed play a significant role in curbing the spread of the virus, and help to eliminate it from the human population. However, scientific efforts to address this challenge are only just beginning. Much remains to be learnt about the virus, its biological properties, epidemiology, etc. At this early stage, there is also a lack of information about specific immune responses against SARS-CoV-2, which presents a challenge for vaccine development.

This study has sought to assist with the initial phase of vaccine development, by providing recommendations of epitopes that may potentially be considered for incorporation in subunit vaccine designs. Despite having limited understanding of how the human immune system responds naturally to SARS-CoV-2, these epitopes are motivated by their responses that they have recorded in SARS-CoV (or, for the case of T cell epitopes, to at least confer MHC binding), and the fact that they map identically to SARS-CoV-2, based on the available sequence data (as of 9 February 2020). The identical map between SARS-CoV and SARS-CoV-2, and the absence of any mutation in the identified epitopes among the available SARS-CoV-2 sequences (as of 9 February 2020), suggests their potential for eliciting a robust T cell or antibody response in SARS-CoV-2.

Research efforts directed towards the design and development of vaccines for SARS-CoV-2 are increasing, and some related analyses are already being reported in distinct, parallel studies. Notably, a preliminary analysis of linear B cell epitopes has been reported online on the ViPR database website (https://www.viprbrc.org/brcDocs/documents/announcements/Corona/2019-nCoV-ViPR-report_24JAN2020.pdf). Different from our study, which is focused on the linear and discontinuous SARS-CoV-derived epitopes, that analysis considered linear B cell epitope data for all Betacoronaviruses from human hosts. While only a summary of the results has been provided so far, preventing direct comparison of the individual epitopes, the number of linear B cell epitopes reported to map identically to SARS-CoV-2 is comparable to our findings.

A recent study has also predicted T cell epitopes for SARS-CoV-2 that may be presented by a population from the Asia-Pacific region (Ramaiah & Arumugaswami, 2020). Again, there are multiple differences to our work. First, the focus of that study was on MHC Class II epitopes, while here we considered both MHC Class I and II epitopes. Interestingly, while we found a few MHC Class II epitopes using our approach (Table S3), only one of these (HLA-DRB1*01:01) appeared in our identified epitope set (Table 3), due to their comparatively low estimated population coverage. Second, computational tools were used to predict MHC Class II epitopes in (Ramaiah & Arumugaswami, 2020), while here we analysed the SARS-CoV-derived epitopes that have been determined experimentally, using either positive T cell or MHC binding assays, and which match identically with the available SARS-CoV-2 sequences (as of 9 February 2020).

We acknowledge that this is a preliminary analysis based on the limited sequence data available for SARS-CoV-2 (as of 9 February 2020). As the virus continues to evolve and as more data is collected, it is expected that additional mutations will be observed. Such mutations will not affect our analysis, provided that they occur outside of the identified epitope regions. If mutations do occur within epitope regions, then these epitopes may be further screened in line with the conservative filtering principle that we have employed, thereby producing a more refined epitope set.

Further experimental studies (T cell and B cell assays) are required to determine the potential of the identified epitopes to induce a positive immune response against SARS-CoV-2. This would help to further refine the reported epitope set, based on observed immunogenicity; an important consideration for immunogen design. Overall, as the identified set of epitopes map identically to SARS-CoV-2, they present potentially useful candidates for guiding experimental efforts towards developing universal vaccines against SARS-CoV-2.

## Data and code availability

All sequence and immunological data, and all scripts for reproducing the results are available at https://github.com/faraz107/2019-nCoV-T-Cell-Vaccine-Candidates.

## Acknowledgements

We thank all the authors, the originating and submitting laboratories (listed in Supplementary Table 7) for their sequence and metadata shared through GISAID, on which this research is based.

M.R.M. and A.A.Q. were supported by the General Research Fund of the Hong Kong Research Grants Council (RGC) [Grant No. 16204519]. S.F.A. was supported by the Hong Kong Ph.D. Fellowship Scheme (HKPFS).

## Supplementary Figures

**Supplementary Figure 1.**
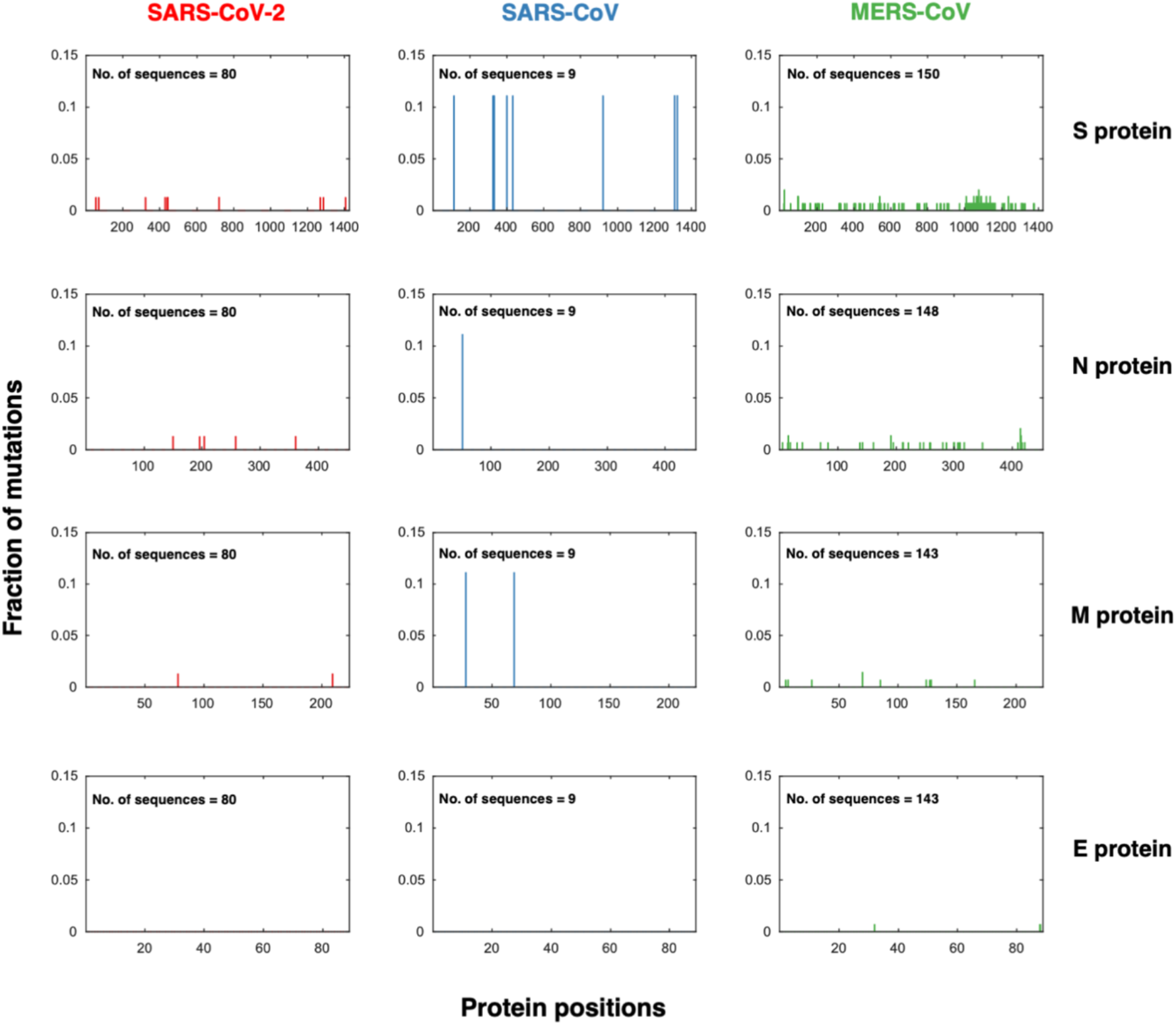
Fraction of mutations in the observed sequences of the structural proteins of the three coronaviruses. Mutation is defined here as an amino acid difference from the reference sequence of the respective coronavirus; accession IDs: NC_045512.2 (SARS-CoV-2), NC_004718.3 (SARS-CoV), and NC_019843.3 (MERS-CoV).

**Supplementary Figure 2.**
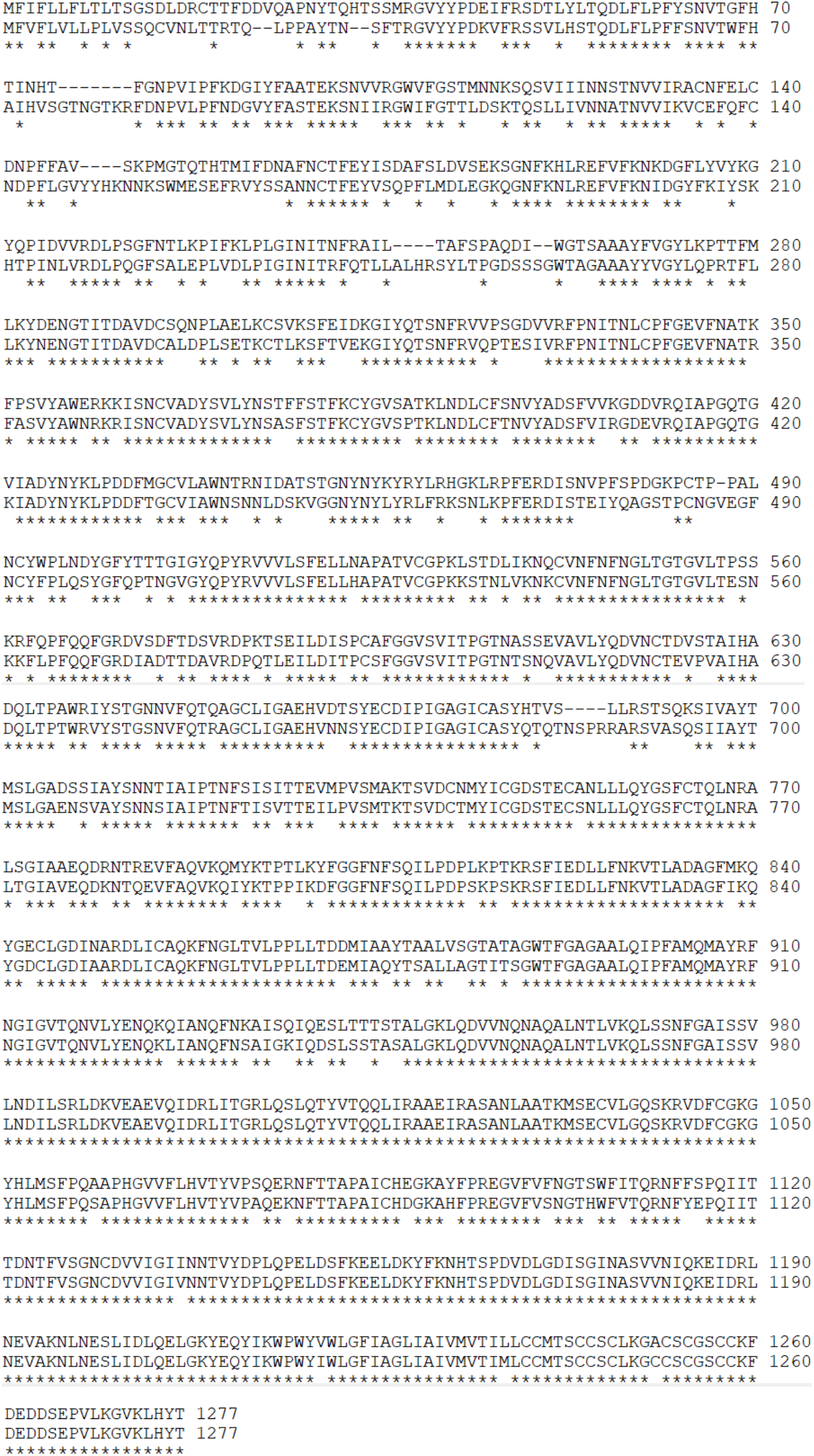
Pairwise sequence alignment of the reference sequences of the S proteins of SARS-CoV and SARS-CoV-2 (accession ID: NP_828851.1 and YP_009724390.1, respectively). Identical residues are indicated by *.

## Supplementary Tables

Table S1: List of GISAID accession IDs for 80 genomic sequences of SARS-CoV-2.

Table S2: List of all SARS-CoV-derived T cell epitopes determined using positive MHC binding assays (with associated MHC allele information available at 4-digit resolution) and found to be identical in SARS-CoV-2.

Table S3: Distribution of all SARS-CoV-derived T cell epitopes obtained using positive MHC binding assays (with associated MHC allele information available at 4-digit resolution) and that are identical in SARS-CoV-2.

Table S4: Set of the SARS-CoV-derived S and N protein T cell epitopes (obtained using positive MHC binding assays) that are identical in SARS-CoV-2 and that maximize estimated population coverage in China (86 distinct epitopes).

Table S5: Estimated global and Chinese population coverages for the individual SARS-CoV-derived S or N protein T cell epitopes (obtained using positive MHC binding assays) that are identical in SARS-CoV-2.

Table S6: SARS-CoV-derived linear B cell epitopes, excluding those in S and N proteins, that are identical in SARS-CoV-2.

Table S7: Acknowledgment table.

